# Consequences of training data composition for deep learning models in single-cell biology

**DOI:** 10.1101/2025.02.19.639127

**Authors:** Ajay Nadig, Akshaya Thoutam, Madeline Hughes, Anay Gupta, Andrew W. Navia, Nicolo Fusi, Srivatsan Raghavan, Peter S. Winter, Ava P. Amini, Lorin Crawford

## Abstract

Foundation models for single-cell transcriptomics have the potential to augment (or replace) purpose-built tools for a variety of common analyses, especially when data are sparse. Recent work with large language models has shown that training data composition greatly shapes performance; however, to date, single-cell foundation models have ignored this aspect, opting instead to train on the largest possible corpus. We systematically investigate the consequences of training dataset composition on the behavior of deep learning models of single-cell transcriptomics, focusing on human hematopoiesis as a tractable model system and including cells from adult and developing tissues, disease states, and perturbation atlases. We find that (1) these models generalize poorly to unseen cell types, (2) adding malignant cells to a healthy cell training corpus does not necessarily improve modeling of unseen malignant cells, and (3) including an embryonic stem cell differentiation atlas during training improves performance on out-of-distribution tasks. Our results emphasize the importance of diverse training data and suggest strategies to optimize future single-cell foundation models.

## Introduction

Intercellular variability in gene expression gives rise to a wide variety of complex biological functions, which are crucial for defining cell types and states [1]. Robustly characterizing this variation across different contexts is a central goal for all areas of eukaryotic biology. In the context of single-cell transcriptomic profiling [2], this goal has been pursued with purpose-driven computational tools, including those intended for cell-type annotation [3], batch integration [4], and perturbation prediction [5]. However, these methods often require many human-in-the-loop steps such as hyperparameter tuning and specification of an appropriate reference, making them difficult to scale to large studies and generalize to unseen datasets. Recently, foundation models have sought to address this problem by pre-training neural network models on large data corpora [6]. These models enable downstream tasks to be performed either directly with embeddings generated from the pre-trained model (i.e., “zero-shot” and “few-shot” evaluation) or by fine-tuning on a subset of new data.

Several foundation models have been developed for single-cell transcriptomics [7–9], drawing inspiration from large language models (LLMs) such as GPT-4 [10]. These models share key features, including (1) transformer-based architectures with attention mechanisms, (2) very large numbers of parameters (commonly on the order of hundreds of millions to billions [11]), and (3) extremely large training datasets, often consisting of tens of millions to hundreds of millions of cells. However, the success of foundation models does not solely depend on model architecture and dataset size. Another critically important factor is training data composition. LLMs are often trained on data from diverse classes (e.g., code, websites, internet articles, and books). To date, research on training data composition for LLMs has included work on restricting data to high-quality sources [12], approaches to infer optimal class weights [13, 14], and algorithms for online optimization of training data composition concurrent with model training [15]. These efforts, among others, have demonstrated how “data mixing” significantly shapes foundation model performance [16]. In contrast, single-cell transcriptomics has operated under the implicit assumption that the largest training corpora will produce the best performing models. However, it has been observed that simply scaling up training datasets does not necessarily yield large gains in model performance [17].

In this work, we investigate how training data composition affects the performance of single-cell foundation models, motivated by three fundamental questions:

1. What is the target distribution of training data for single-cell foundation models, and can we evaluate how well a training corpus represents this target distribution?
2. What is the analog of data mixes (e.g., training text classes) in single-cell data?
3. Given a baseline training corpus, how does the addition of categorically different training data change model performance?

The answers to these questions could have substantial implications for improving training efficiency and model performance. We use the *hierarchy of cell states in development* as a framework for considering diversity in single-cell training data and explore these questions within a tractable biological system, human hematopoiesis (**Figure 1A**). We then use this approach to create various corpora with which to train a deep learning architecture that supports interpretable *post hoc* analysis [18], and we verify our observations in a much larger transformer-based model architecture that more closely matches modern state-of-the-art models [7]. Through this study, we identify three key patterns relating training data composition to model performance and analyze the mechanisms underlying these patterns. First, we show systematically that deep learning methods struggle to generalize to cell types that are not represented in the training data. Next, we demonstrate that including malignant cells in a training corpus does not necessarily improve predictions for cells from unseen cancer subtypes or disease states. Lastly, we show that directed differentiation atlases can provide a source of training data with enough coverage of diverse cell types to enhance performance on out-of-distribution tasks.

**Figure 1.**
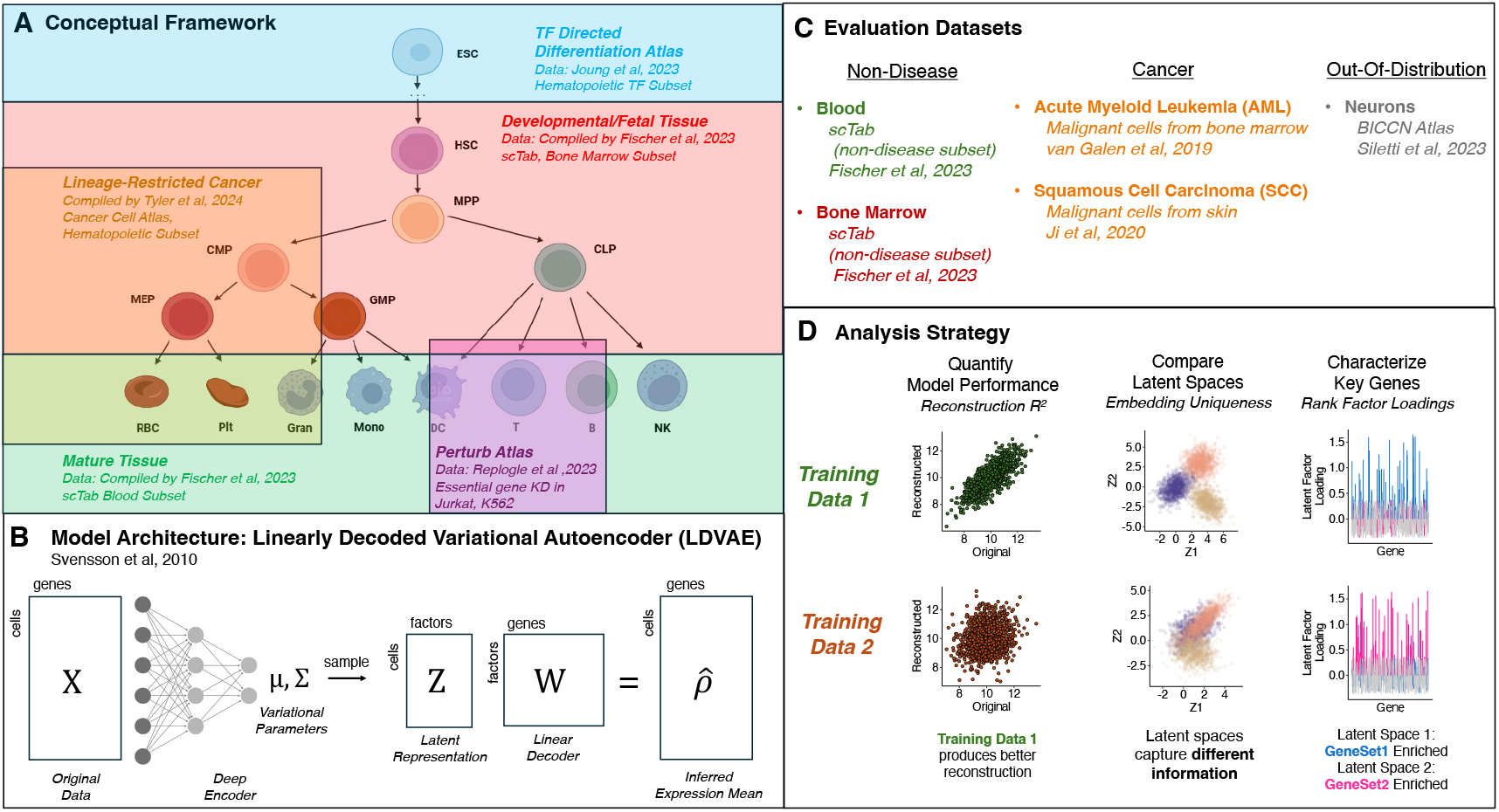
Assessing the influence of training data composition on model performance in single-cell biology. **(A)** Schematic of a hierarchical framework for understanding the distribution of cellular states. Highlighted are different hematopoietic atlases and datasets from the literature and the representative cellular states that they include. **(B)** Schematic of the linearly decoded variational autoencoder model (LDVAE) that is used for the primary analysis. **(C)** Description of the different datasets used for downstream evaluation. **(D)** Description of the specific analytic strategies for evaluating how the composition of training data influences models. These analyses include (i) comparing predictive performance (via reconstruction *R*^2^), (ii) comparing the latent spaces between models trained on different datasets, and (iii) characterizing the key genes in the factor loadings of different models.

## Results

### Overview of analyses and experimental design

We hypothesized that training corpora for single-cell foundation models should – ideally – capture the distribution of possible cellular states. One framework for understanding this distribution is through the *hierarchy of cell states in development*, which connects embryonic cells to mature adult cells through a branching sequence of progressively differentiated progenitors [19]. This framework captures the distribution of observed cellular states by describing the mechanistic processes that gives rise to them. Just as in genetics where phylogenies capture inter-individual variability empowering diverse downstream applications [20], the cellular hierarchy can organize our understanding of inter-cellular variability. This idea is central in developmental biology [21], has guided the engineering of cell state transitions [22], and underpins a leading theory of malignant transformation [23]. More than a century of developmental biology has mapped these hierarchies across human tissues.

In alignment with this view, fitting a “complete” foundation model of single-cell transcriptomic states requires a training corpus that encompasses the entire developmental hierarchy. To illustrate, we considered the well-characterized hematopoietic lineage (**Figure 1A**), which connects central hematopoietic stem-cells (HSCs) to peripheral mature cell types, such as erythrocytes and T cells, through a series of intermediate progenitors [24]. We posited that, while different sources of training data cover distinct subregions of this hierarchy, collectively they would tessellate the entire distributional space. Mature adult tissues (e.g., peripheral blood cells) span the leaves of the tree (**Figure 1A** in green). Developmental tissues (e.g., bone marrow cells) cover progenitors that give rise to the mature adult cells (**Figure 1A** in red). Deranged cellular states (e.g., cells from malignancies) diverge from typical states but may retain some lineage restriction (**Figure 1A** in orange). Perturbation atlases (e.g., cells from massive Perturb-seq experiments) explore local microstates surrounding particular nodes of the tree (**Figure 1A** in purple). And, lastly, directed differentiation atlases (e.g., cells from massively parallel transcription factor upregulation experiments in embryonic cells) explore very early progenitors, including those that potentially cannot be sampled in an adult organism (**Figure 1A** in blue).

We trained representation learning models with these data sources individually and in combination, and then evaluated their predictive performance on “in-distribution” and “out-of-distribution” datasets to identify strengths and weaknesses of different training corpus construction strategies (**Methods**). Briefly, we first trained a “Blood (Baseline)” model exclusively on ca. 100k healthy (101,879 cells exactly) peripheral blood cells [25]. Next, we implemented individual “BoneMarrow” [25], “BloodCancer” [26], “PerturbSeq” [27, 28], and “TFAtlas” [29] models, each trained on ca. 100k cells (101,879 exactly) exclusively from their respective studies. Lastly, we fit “Blood+” combination models, all trained on ca. 200k cells (203,758 exactly) with half of the cells being healthy peripheral blood cells and the other half coming from one of other sources. For these experiments, we used a linearly decoded variational autoencoder (LDVAE) [18] (**Methods**), which is based on the single-cell variational inference (scVI) framework [30] that has been shown to be competitive with transformer-based models equipped with many more parameters [9, 31]. The advantage of the LDVAE model is that it uses a linear transformation instead of a neural network as a decoder. This enhances its interpretability because it enables the direct estimation of how individual genes contribute to variables learned in the latent space.

To understand the impact of each data type, we computed the zero-shot reconstruction accuracy (via R-squared, *R*^2^) for each trained LDVAE model across five different evaluation datasets containing blood cells, bone marrow cells, acute myeloid leukemia (AML) malignant cells, squamous cell carcinoma (SCC) malignant cells, and neuronal cells (**Methods**). These evaluation datasets were selected to represent a spectrum of in-versus out-of-distribution settings with respect to the training corpora considered. We focused on the reconstruction task because having the ability to represent and reconstruct cellular states is fundamental to many downstream single-cell applications. Importantly, this evaluation does not rely on labels (such as cell type or perturbation status), which may not always be available or inconsistently defined across datasets. This collective analysis resulted in 225 model comparisons, each being robustly reproduced across five different random seeds (**Figure S1**).

For our Blood (Baseline) model, we observed that the reconstruction accuracy was greatest in held-out peripheral blood cells (*R*^2^ = 0.69), worse in bone marrow and hematological cancer (AML) cells (*R*^2^ = 0.38 and 0.33, respectively), worse still in non-hematological cancers (SCC) (*R*^2^ = 0.24), and very poor in neurons (*R*^2^ = 0), an expected pattern of results given its training corpus (**Figure 2**). Changes to the training data composition revealed a more complex pattern of results. Here, we identified three distinct patterns that both capture variability in performance across models and have implications for training future ones. First, model performance decreases when evaluation datasets include cell types not represented in the training data. Second, including cancer data in the training corpus does not improve the reconstruction of unseen cancer data. Third, training with data from a directed differentiation atlas (i.e., activation of many transcription factors in embryonic stem cells) improves performance for out-of-distribution cell types. In the following sections, we describe our in-depth empirical analyses to generate hypotheses as to why these patterns occur in practice.

**Figure 2.**
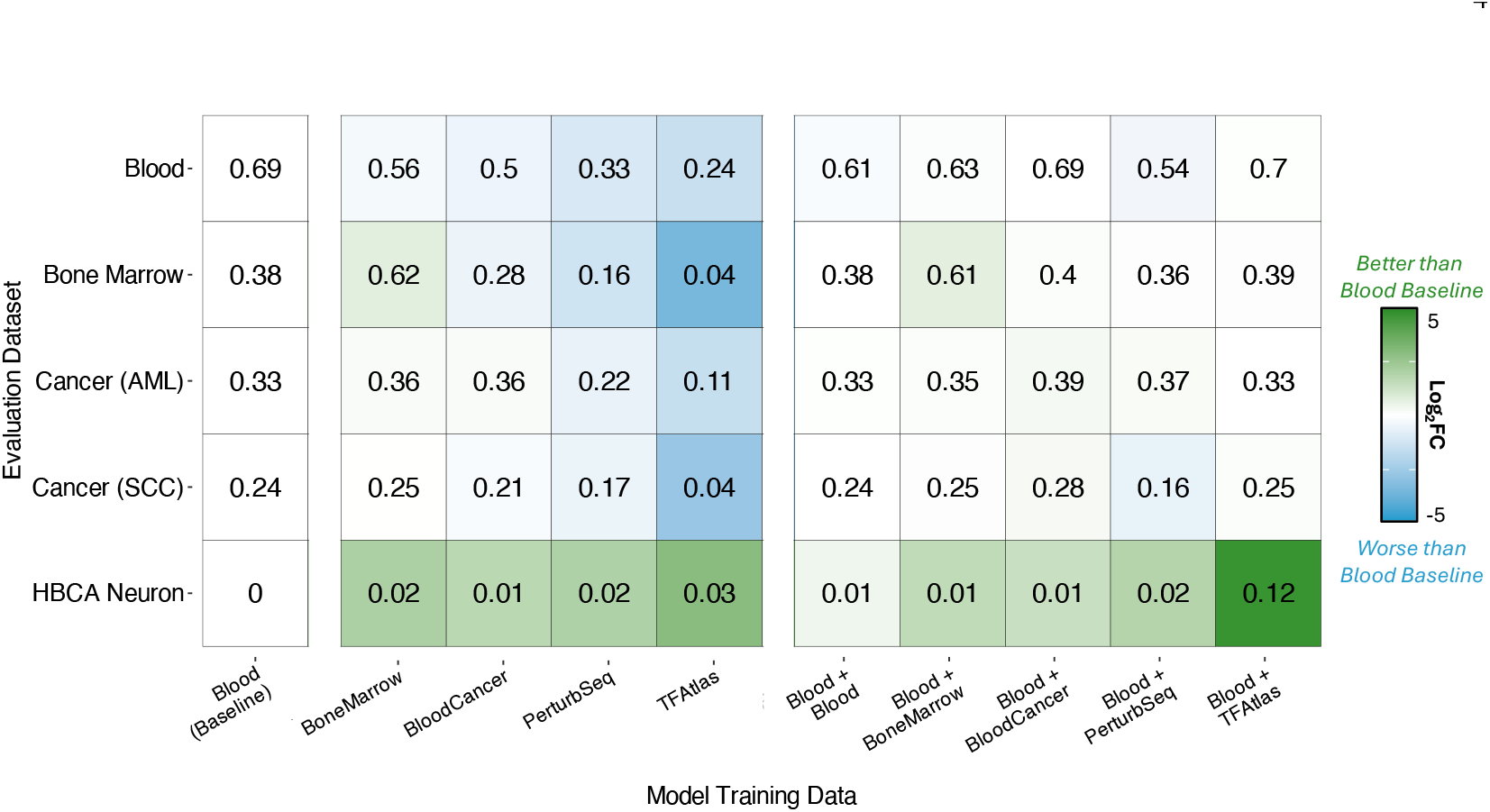
Estimates of reconstruction accuracy for various training and evaluation datasets using LDVAE. Different training data compositions are listed across each column. Here, the “Blood (Baseline)” model is trained exclusively on ca. 100k healthy peripheral blood cells [25]. The “BoneMarrow” [25], “BloodCancer” [26], “PerturbSeq” [27, 28], and “TFAtlas” [29] models are also trained on ca. 100k cells coming exclusively from their respective studies. The “Blood+” combination models are all trained on ca. 200k cells, with half being healthy peripheral blood cells and the other half being from the other source. Numerical results in the heatmap are based on the average reconstruction accuracy (Pearson’s *R*^2^) for highly-variable genes across each of the evaluation datasets shown across the rows. Averages are based on five different random seeds for each analysis. Within a given row, the color of each cell indicates the log_2_ fold change of the reconstruction *R*^2^ relative to the “Blood (Baseline)” model (with darker green representing higher log_2_ fold changes). See **Methods** for a detailed description of each training and evaluation dataset.

### Deep learning models generalize poorly to unseen cell types

Our analysis revealed that reconstruction accuracies were consistently lower when models were evaluated on datasets containing unseen cell types and/or states (**Figure 2**). For example, a model trained on peripheral blood cells performed poorly on bone marrow cells (*R*^2^ = 0.38) compared to a model trained on bone marrow cells (*R*^2^ = 0.62). Similarly, a model trained on bone marrow cells performed worse on peripheral blood cells (*R*^2^ = 0.56) than a model actually trained on peripheral blood cells (*R*^2^ = 0.69). All models performed poorly on neurons (*R*^2^ less than 0.15), which were not included in any of the training datasets.

Interpreting these observations is confounded by the fact that each training dataset contains a mixture of many cell types, with complex relationships between cell types. To disentangle this effect, we implemented two additional models trained exclusively on peripheral blood but holding out a single cell type, either monocytes (“No-Mono”) or B cells (“No-Bcell”) (**Figure 3A**; **Methods**). The No-Mono and No-Bcell models demonstrated worse reconstruction accuracy for monocytes and B cells, respectively, compared to a model trained on all blood cell types (**Figure 3B**). To understand the source of this decreased performance, we examined the prediction residuals from the reconstructions of the No-Mono and No-Bcell models (**Methods**). The distributions of these residuals were extremely heavy-tailed, indicating that a small subset of genes was reconstructed very poorly (**Figure 3C**). These genes were immediately recognizable as having key functions in the held-out evaluation cell type. For example, *FTL*, a gene highly expressed in monocytes compared to other peripheral blood mononuclear cells (PBMCs) [32], was poorly reconstructed in the No-Mono model, while *CD74*, a key regulator of B cell maturation [33], was poorly reconstructed in the No-Bcell model (**Figure 3C**).

**Figure 3.**
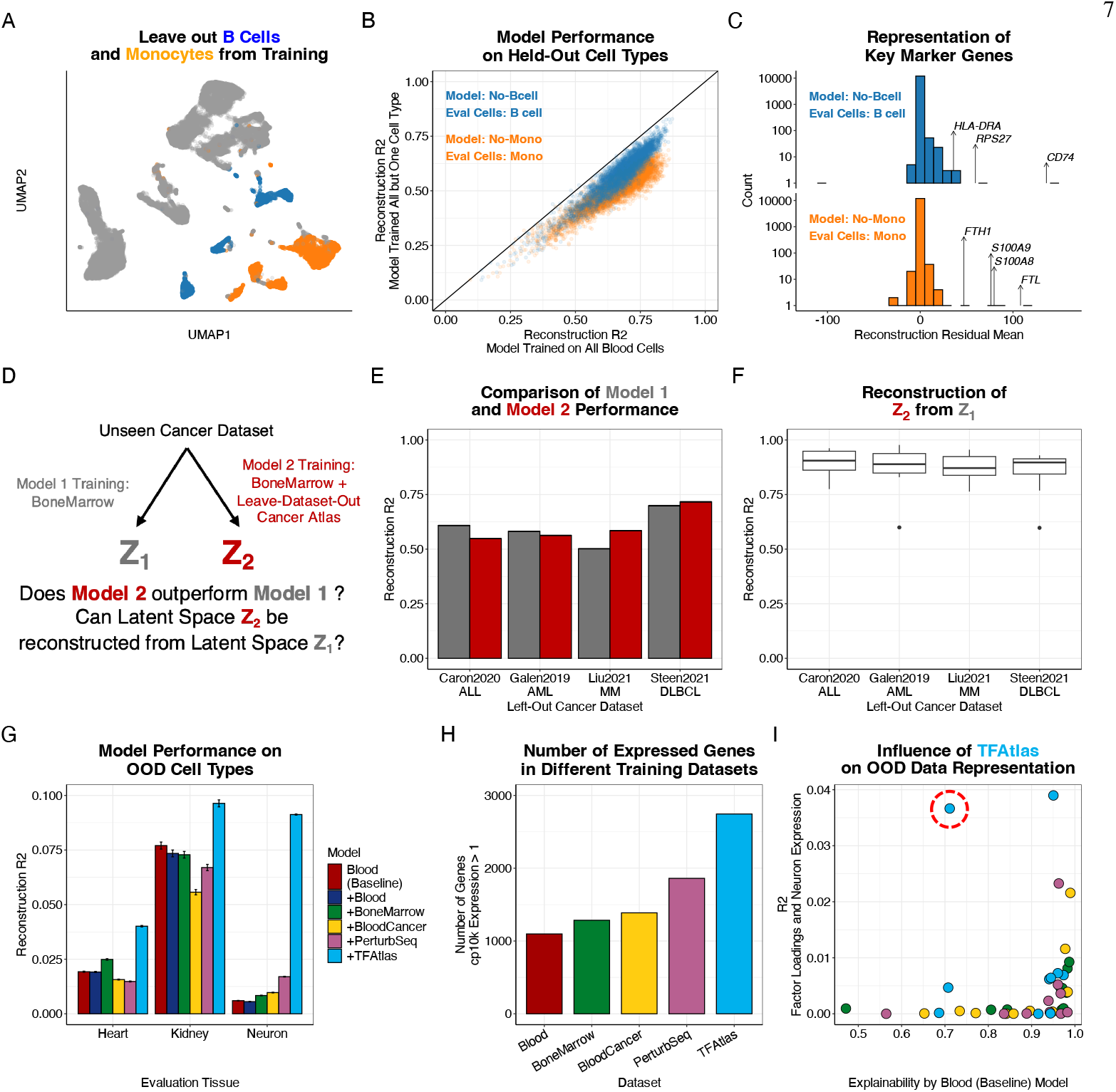
Explaining differences in model performance across training datasets. **(A)** Uniform manifold approximation and projection (UMAP) showing the Blood (Baseline) dataset, with the two cell types (B cells and monocytes) highlighted (blue and orange, respectively) to be held-out during training. Visualized are all 101,879 blood cells from the Blood (Baseline) training dataset. Of these, there are 16,096 monocytes (orange) and 9,723 B cells (blue). **(B)** Reconstruction accuracy on a particular cell type for a model trained on all blood cells versus a model trained with that cell type held-out (blue, B cells, *N* = 3651; orange, monocytes, *N* = 6052). We refer to the model holding-out monocytes as “No-Mono” and the model holding-out B cells as “No-Bcell”. **(C)** Visualization of gene-wise reconstruction residuals for cross-cell type model evaluations: (top) train a model with B cells and test on monocytes, and (bottom) train a model with monocytes and test on B cells. **(D)** Schematic of an analytic strategy to assess the impact of adding cancer data to the BoneMarrow training corpus. The idea is that if the cancer data is adding new information, then the latent space from a BoneMarrow-only model should struggle to explain the variance in the latent space generated by a BoneMarrow+Cancer model. **(E)** Visualization of the reconstruction accuracy (y-axis) for various held-out cancer datasets (x-axis) comparing models trained on bone marrow only cells (grey) versus bone marrow and other cancer cells (red). Results are based on five different seeds, with error bars representing the standard errors across replicates. **(F)** Visualization of *R*^2^ indicating how well the latent space of the BoneMarrow+Cancer model can be reconstructed from the BoneMarrow only model latent space. There are *K* = 10 latent dimensions per boxplot. The center line indicates the median, hinges indicate first and third quartiles, and the whiskers indicate largest value no more than 1.5 × IQR (interquartile range) away from hinge. Points beyond these whiskers are plotted individually. (**Methods**). **(G)** Visualization of reconstruction *R*^2^ for each “Blood+X” model (where X is Blood, BoneMarrow, Cancer, PerturbSeq, or TFAtlas) on three out-of-distribution datasets. Teal bars indicate estimates from the Blood+TFAtlas model. Error bar indicates standard error of mean across cells in the evaluation. There are *N* = 9,926 cells in the Neuron dataset, *N* = 13,571 in the Heart dataset, and *N* = 8,468 in the Kidney dataset. **(H)** Visualization of the number of genes with cp10k expression > 1 in each training dataset. **(I)** The x-axis visualizes the ability to reconstruct the latent space of the Blood+ models from the latent space of the Blood (Baseline) model. The y-axis shows the correlation between latent variable gene-wise loadings and the mean expression level in neurons. Teal points indicate estimates from the Blood+TFAtlas model, while the other points represent latent variables from the other Blood+ models. The red circle highlights a latent factor from the Blood+TFAtlas model that is poorly explained by the Blood (Baseline) model and correlated with neuronal expression.

Lastly, while perturbation data has been hypothesized to facilitate out-of-distribution inference [34], we observed no significant improvement in the ability to generalize to unseen cell types for models trained with Perturb-seq data from erythroid- and leukocyte-derived cell lines (**Figure 2**). For example, a model trained on peripheral blood cells performed better on bone marrow cells (*R*^2^ = 0.38) than a model trained only on cells from Perturb-seq experiments (*R*^2^ = 0.16). Moreover, training a model on a dataset that combined the blood and perturbed cells did not yield any additional improvement (*R*^2^ = 0.36).

Expanding on recent analyses of out-of-distribution performance [5, 35], these findings suggest that models trained on an incomplete corpus fail to generalize because they cannot adequately represent key genes that define unseen cell types.

### Including malignant cells in the training corpus has a modest impact on cancer data evaluations

Modeling data from human malignancies is a critical target application for single-cell foundation models [8]. Such models could potentially map the distribution of cellular states within individual tumors and identify patient-specific treatment strategies. The held-out evaluations for models trained on only healthy cells (**Figure 3A-C**) suggested that incorporating data from human malignancies could improve the ability to represent broader sets of cellular states (e.g., disease phenotypes). However, including a large atlas of diverse hematological cancer data did not significantly improve performance on a held-out acute myeloid leukemia (AML) evaluation dataset or on a held-out, non-hematological squamous cell carcinoma (SCC) dataset (third and fourth rows in **Figure 2**).

To investigate this pattern further, we trained multiple LDVAEs using healthy bone marrow and hematological cancer atlas data, systematically leaving out one of four datasets from the hematological cancer atlas (**Figure 3D**; **Methods**). We then evaluated each model’s performance on the held-out dataset and compared it to a baseline model trained exclusively on healthy bone marrow data. Across all leave-one-dataset-out models, the inclusion of cancer data generally had only modest effects on predictive performance compared to a model trained only on healthy bone marrow data (**Figure 3E**). The one exception to this trend was the model trained on the hematological cancer atlas excluding the large multiple myeloma dataset from Liu et al. [36], which produced a 17% increase in reconstruction accuracy relative to training on only healthy bone marrow data. This is likely due to the fact that other studies in the cancer atlas included multiple myeloma cells [26], whereas the other malignancies were not as well-represented when they were held-out during training.

We hypothesized that the modest changes in performance could be due to the fact that cancer cells, while on average occupying distinct states, share axes of variation already represented within the diversity of healthy cells (e.g., non-clinical variability in proliferation across healthy cells). To test this hypothesis, we attempted to reconstruct the latent space of models trained with cancer data from the latent space of models trained only on healthy bone marrow data (**Methods**). In all models trained with leaving out one dataset, this reconstruction was successful, with linear regression producing mean *R*^2^ values exceeding 0.85 (**Figure 3F**). For example, in a model trained with both bone marrow and cancer data excluding cells from the Caron et al. [37] acute lymphoblastic leukemia study, one latent dimension was found to be enriched for genes involved in mitosis (including *TOP2A, PLK1, CDK1*, and *CENPA*) (**Figure S2**). While these features are likely relevant to malignant pathology, the variance in this dimension was still explained accurately by the latent space of a model trained only on bone marrow data (*R*^2^ = 0.96; **Figure 3F**). However, we also note that inclusion of data from a matched malignancy in training data, as in our multiple myeloma experiment, improved model performance (**Figure 3F**). Altogether, these findings suggest that the variability introduced by cells in this particular cancer atlas do not augment the learned latent space with generalizable axes that can be used to model out-of-distribution malignancies.

### Integrating transcription factor atlases into training corpora improves out-of-distribution performance

Since including both perturbation and cancer data into the training corpora had only a small impact on the out-of-distribution performance for the models we trained, we next sought to evaluate the impact of directed differentiation atlases (**Figure 1A** in blue). Systematic activation of transcription factors in embryonic stem cells can generate a diverse range of early progenitor cells [29]. We hypothesized that models trained on such data could leverage this variability to improve performance on unseen cell types. Notably, training with a directed differentiation atlas (“TFAtlas”) yielded the greatest improvements not in developing tissue (i.e., bone marrow), but in mature neurons (**Figure 2**). While the absolute reconstruction accuracy remained limited, there was a significant relative gain in prediction performance. This result is important given two factors: (1) our previous findings of poor generalization to unseen cell types, as the TFAtlas does not include true neurons, and (2) the fact that the TFAtlas dataset was subsetted to enrich for perturbations of blood-expressed transcription factors (**Methods**).

To assess the robustness of this finding, we repeated our analyses using two additional unseen, out-of-distribution cell types: heart and kidney cells (**Methods**). Consistent with the results observed in neurons, the Blood+TFAtlas model outperformed other models within these populations (**Figure 3G**). We hypothesized that the TFAtlas enhances the model’s ability to generalize out-of distribution because the TFAtlas captures non-zero expression across more genes, broadening the model’s applicability. Indeed, the TFAtlas contains the highest number of substantially expressed genes (where substantial expression is cp10k > 1; **Figure 3H**) among all the training datasets used in our analyses. We note that this large number of expressed genes is likely due to a combination of (1) the “permissive” expression state of embryonic stem cells at baseline [38], and (2) the diversity of expression states generated by TF perturbations. The latent space of the Blood+TFAtlas model should thus include features that cannot be explained from the latent space of the Blood (Baseline) model. Here, we observed a latent dimension in the Blood+TFAtlas model (red circled point in **Figure 3I**) that could not be reconstructed from the Blood (Baseline) model (**Methods**) and was correlated with average gene expression of neuronal cells (**Figure S3**). Many of the top-loaded genes for this latent dimension were enriched for neuronal functions, especially synaptic function (e.g., *CSMD2, RIMS2*, and *DLG2*). Notably, one of the top-loaded genes was a subunit of the glutamatergic kainate-type receptor, *GRIK3*. These findings indicate that, while the TFAtlas data does not contain actual neurons, it does include axes of intercellular variability that are associated with neuronal gene expression. In turn, the presence of such a correlated latent variable was associated with improved out-of-distribution performance on the unseen neuronal evaluation dataset.

### Training data composition drives predictive performance regardless of model architecture complexity

Our experiments thus far utilized (1) the LDVAE framework, which has a less sophisticated architecture than recent transformer-based models [7–9] and (2) a baseline model trained exclusively on cells from a single tissue (whole blood), rather than one trained on a diverse cellular atlas. To explore the impact of these parameters, we conducted additional analyses by (1) training Geneformer models [7] instead of LDVAEs, and (2) using an identically-sized (*N* = 101,879) subsample of the entire scTab dataset (rather than blood alone) as the baseline training corpus. This resulted in a 2 × 2 array of experiments for comparison: an LDVAE model trained on the Blood (Baseline) dataset, an LDVAE model trained on an Atlas (Baseline) dataset, a Geneformer model trained on the Blood (Baseline) dataset, and a Geneformer model trained on an Atlas (Baseline) dataset.

We first evaluated the performance of the Geneformer Blood (Baseline) models (**Figure S4**). Since Geneformer produces ranked gene expression outputs rather than fully reconstructed expression estimates across all genes, we computed the correlation between ranks for each training/evaluation combination [31]. The results mirrored those of the LDVAE Blood (Baseline) models (**Figure S5**) where the worst performance was observed for out-of-distribution cell types. For example, the Geneformer Blood (Baseline) model had a mean rank correlation of 0.62 when being evaluated on a dataset with blood cells; however, it only had a mean rank correlation of 0.06 on an evaluation dataset with neurons (**Figure S4**). Similarly, the addition of malignant cells in the training dataset did not significantly improve performance of Geneformer on the cancer evaluation datasets (e.g., the Cancer AML evaluation yielded a mean rank correlation of 0.87 for the Blood+Blood model and 0.91 for the Blood+BloodCancer model). Notably, however, the HBCA-Neuron evaluation results differed between the LDVAE and Geneformer Blood (Baseline) models (**Figure S5**). While neurons remained the worst-performing evaluation set for the Geneformer models, we did not observe large performance improvements from the inclusion of the TFAtlas data in the same way that we saw a significant increase in the LDVAE analysis. We speculate that this difference may be due to the normalization and tokenization procedures of Geneformer. These procedures include normalizing across cells, compressing expression data into ranks, and limiting to highly expressed genes, whereas in contrast, LDVAEs are models for transcriptome-wide observed expression counts.

Next, we evaluated the LDVAE and Geneformer Atlas (Baseline) models (**Figure S6-S9**). Expanding the baseline training corpus to include a larger atlas had the greatest impact on the neuron evaluations (**Figure S6**), likely because the neuron dataset was no longer out-of-distribution due to the presence of brain cells in the training dataset. Consequently, the contribution of the TFAtlas was diminished (**Figure S7** and **S8**), suggesting that the perturbed embryonic stem cells with neuron-like expression added little value beyond the true neurons already observed during training. For the cancer evaluations, the pattern of log_2_ fold changes was nearly identical when comparing the LDVAE Atlas (Baseline) to the LDVAE Blood (Baseline) models (correlation for the AML evaluation = 0.97, correlation for the SCC evaluation = 0.80; **Figure S9**), with slopes also not significantly different than 1 (slope for the AML evaluation = 0.89 with *P* = 0.2 based on a t-test under a null hypothesis that the slope = 1; slope for the SCC evaluation = 1.18 with *P* = 0.52 based on a t-test under a null hypothesis that the slope = 1). The trends observed with the LDVAE architecture were similar to trends observed with the Geneformer architecture when comparing the Blood (Baseline) to the Atlas (Baseline) (**Figure S6**).

These findings highlight that while varying model architecture can influence performance, the patterns of generalization, particularly for out-of-distribution contexts, remain constrained by the composition and diversity of the training corpus.

## Discussion

Foundation models have the potential to unify and transform many areas of single-cell biology. In this study, we investigated how training data composition influences the performance of deep learning models of single-cell transcriptomics, using the human hematopoietic system as a framework to guide investigation. Our findings illuminate key factors impacting out-of-distribution generalization, the relative contributions of normal versus pathological cells in training, and the unexpected benefits of directed differentiation atlases. These results establish a starting point for developing rational principles to guide the design of training corpora for single-cell foundation models. We anticipate that further exploration of these principles will unlock substantial improvements in model performance.

To ensure tractable inference, we constrained our analyses to specific datasets, model architectures, and evaluations. However, these choices introduce limitations. For example, the principles identified in this work may vary in applicability across tissue compartments (i.e., outside of hematopoiesis), species (i.e., mice or non-human primates), or disease states (i.e., non-malignant disease states). Future research should extend these analyses to other biological contexts. Nonetheless, current training datasets for single-cell foundation models are heavily biased towards blood cells [7–9], suggesting that the patterns we observed here are likely reflective of current state-of-the-art deep learning methods. Furthermore, while our focus on reconstruction accuracy provides a foundation for representing cellular states, it raises questions about whether similar patterns would emerge for downstream tasks such as cell-type classification or perturbation prediction [39]. We prioritized reconstruction accuracy because it underpins all downstream tasks (i.e., models must be able to represent cell states before they can estimate complex functions of cell state) and does not require high-quality data labels.

Our findings regarding the representation of malignancies are particularly notable. They suggest that while datasets of healthy tissue lack malignant cells, they still encompass axes of variation that are sufficient to represent (at least some) unseen malignant states. This insight has significant implications for training future single-cell foundation models. For example, adding smaller, heterogeneous cancer datasets (e.g., the Curated Cancer Cell Atlas used here; [26]) to large, systematically collected datasets of healthy cells [40] might introduce more integration challenges (i.e., batch effects) than performance benefits. While we caution that this hypothesis has not been exhaustively validated for malignancies—and certainly not for the broader spectrum of human pathologies—our results underscore the need for further research into the costs and benefits of including pathology-derived data into foundation model training corpora.

Our results also highlight the unique value of directed differentiation atlases in improving out-of-distribution performance. While these datasets were originally designed to systematically study the individual and combinatorial effects of human transcription factors [29], our analyses point to a second use of such differentiation techniques: to generate single-cell data with immense variability in expression states. Such variability enables foundation models to learn generalizable axes of cellular variation. This is particularly valuable because, experimentally, these axes can be observed within a single dataset with minimal technical artifacts. Our findings suggest a synergy between basic research on human transcription factors, new methods in functional genomics, and the engineering of foundation models, where the collection of larger directed differentiation atlases could drive advances across fields.

Lastly, and more broadly, our work emphasizes the importance of training data composition in biological applications of machine learning. While many foundation models continue to prioritize larger datasets and more advanced architectures, our results suggest that performance gains may be achievable through principled optimization of smaller training datasets, mirroring observations in natural language [12]. We call for renewed attention to this critical factor in single-cell biology, as it has the potential to enhance the utility and generalizability of deep learning models in transcriptomics and beyond.

## Methods

### Training datasets

We used five different datasets to create unique corpora for training the deep learning models implemented in this study.

- **scTab [25]:** This dataset is a subset of the CELLxGENE census [41] (version 2023-05-15) and consists of 22.2 million cells from 164 unique cell types, 5,052 unique human donors, and 56 human tissues. Any training corpus labeled with “Blood” or “BoneMarrow” was created by subsampling from blood and bone marrow cell populations within this resource, respectively.
- **Curated Cancer Cell Atlas (3CA) [26]:** This repository consists of 124 different studies spanning over 40 cancer types and 2,822 samples (version 2024-07-30). To create a “BloodCancer” training corpus, we subsetted the hematological datasets in this atlas to those that (1) had integer count data available and (2) were sequenced using either 10x, MARS-Seq, or Seq-Well technologies (note that the other technologies, SmartSeq2 and HiSeq2000, were only used for one dataest each). Using this criteria left us with the following cancer datasets for our study: acute lymphoblastic leukemia from Caron et al. (2020) [37]; multiple myeloma from Liu et al. (2020) [36]; myeloproliferative neoplasm from Nam et al. (2019) [42]; B cell lymphoma from Roider et al. (2020) [43]; diffuse large B cell lymphoma from Steen et al. (2021) [44]; follicular lymphoma from Zhang et al. (2019) [45]; multiple myeloma from Cohen et al. (2021) [46]; and multiple myeloma from Ledergor et al. (2018) [47]. For each of these datasets, we subsampled only cells that were annotated as being “malignant”.
- **K562 [27] and Jurkat [28] Perturb-seq:** We utilized essential gene Perturb-seq datasets (CRISPR-based screens with single-cell RNA-sequencing readouts). Specifically, this included 2,057 CRISPR interference (CRISPRi) perturbations in the K562 cell line [27] and 2,393 CRISPRi perturbations in the Jurkat cell line [28]. The perturbed genes were found to have significant growth effects in the Cancer Dependency Map (DepMap) project [48]. To maximize the expression variability coming from these datasets within our “PerturbSeq” training corpora, we filtered such that we retained only cells that received a targeting guide RNA (i.e., excluding the “control” cells receiving non-targeting guide RNAs).
- **Transcription Factor (TF) Atlas [29]:** This resource contains RNA-sequencing data after the overexpression of one of 3,548 human TF splice isoforms with CRISPR activation (CRISPRa) in human embryonic stem cells. To enrich our “TFAtlas” training dataset for blood-active TFs, we first ranked all perturbed transcription factors by their degree of expression in whole blood bulk sequencing data from the Genotype-Tissue Expression (GTEx) Portal [49]. Specifically, we z-scored (mean centered and unit standardized) the distribution of expression for each TF across tissues, and retained only TFs where the whole blood z-score was greater than -0.25 (i.e., these TFs were deemed to have greater expression in whole blood versus other tissues). The threshold of -0.25 was decided qualitatively based on visual inspection of the distribution of these z-scores, to jointly maximize both specificity and number of cells retained.

For each training dataset, we applied the same series of preprocessing steps. We filtered out cells with a proportion mitochondrial counts greater than 0.1 and a total number of unique molecular identifiers (UMIs) greater than 1000. We then subsetted each dataset to have 101,879 cells, which was the size of the smallest study described above (i.e., the Curated Cancer Cell Atlas). Note that in the text, we say that sample sizes were multiples of 100k for simplicity. For the Blood, BoneMarrow, Perturb-seq, and TFAtlas datasets we performed this subsetting via uniform random downsampling. No subsetting was necessary to create the BloodCancer corpus.

### Evaluation datasets

We used five different datasets to evaluate the reconstruction accuracy of each trained model. Details for each are given below:

- **Blood**: This evaluation dataset consisted of 38,251 blood cells that were held-out from the Blood training corpus, which was derived from the original scTab dataset [25].
- **Bone Marrow:** This evaluation dataset consisted of 12,732 bone marrow cells that were held-out from the BoneMarrow training corpus, which was derived from the original scTab dataset [25].
- **Cancer (AML):** This evaluation dataset consisted of 14,702 acute myeloid leukemia malignant cells from van Galen et al. [50], which can be accessed via the Curated Cancer Cell Atlas [26].
- **Cancer (SCC):** This evaluation dataset consisted of 10,583 squamous cell carcinoma malignant cells from Ji et al. [51], which can be accessed via the Curated Cancer Cell Atlas [26].
- **Human Brain Cell Atlas (HBCA):** This evaluation dataset consisted of a random set of 9,926 neuronal cells subsampled from the Human Brain Cell Atlas [52]. This resource is publicly available online and can be accessed via GitHub at github.com/linnarsson-lab/adult-human-brain.

For each of these datasets, we performed the same filtering for mitochondrial read proportion and number of UMIs as described above for the training datasets. The numbers of cells listed above describe the post quality-controlled sample sizes per dataset.

### Linearly decoded variational autoencoders

In this section, we briefly review the linearly decoded variational autoencoder (LDVAE) framework, which we use as one of the main deep learning models in our study. A complete detailed breakdown of the method can be found in Svensson et al. [18]. Consider a study with single-cell RNA sequencing expression data for *i* = 1, …, *N* cells that each have measurements for *j* = 1, …, *G* genes. Let this dataset be represented by an *N* × *G* matrix **X** where the row-vector **x**_*i*_ = [*x*_*i*1_, …, *x*_*iG*_] denotes the expression profile for the *i*-th cell. We will denote **z** to be a *K*-dimensional vector of stochastic latent variables (i.e, a representation of the single-cell data). A typical variational autoencoder (VAE) [53] has two key components: (1) an encoder which maps the original expression vector of each cell **x**_*i*_ to a lower dimensional space characterized by **z**, and (2) a decoder which takes those learned latent variables **z** and attempts to reconstruct the expression distribution parameters. Intuitively, a successfully trained model aims to minimize the loss between the original and reconstructed data across *N* cells in the training set. The single-cell variational inference (scVI) [30] model is a particular VAE. It assumes the following generative process

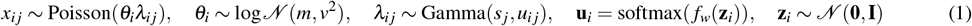

where the expression of the *j*-th gene in the *i*-th cell *x*_*i j*_ is assume to follow a Poisson distribution, *θ*_*i*_ is a log-normal distributed random variable representing the exposure or count depth of a cell characterized by hyperparameters *m* and *v*^2^, *s* _*j*_ represents the overdispersion of a gene, the Gamma distribution is parameterized by its shape and mean, and the *K*-dimensional latent variable **z**_*i*_ follows a standard normal distribution with a zero mean vector and identity covariance matrix. Note that, in our analyses, we refer to the inferred expression mean (**û**_*i*_) as the reconstructed expression (which we denote as 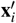). Autoencoders typically utilize neural network architectures *f*_*w*_() to infer the latent variable values **z** (the representation) and to perform reconstruction of the original data **x**_*i*_. However, the representations inferred with VAEs are not directly interpretable. In the LDVAE, the neural network decoder is replaced with a linear function such that

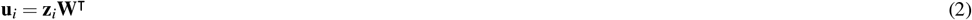

where **W** is an *J* × *K* is a factor loadings matrix effectively detailing how important the *j*-th gene is in describing the *k*-th latent variable, thus providing a direct link between cell representation and gene expression [18] (e.g., **Figures 1D** and **3H**).

### LDVAE implementation details

We used the implementation of LDVAE that is publicly available as part of the `scvi-tools` package (release 1.1.2) [54] under the `LinearSCVI` class. The default parameters were used to set up the architecture of these models. Specifically, the number of nodes per hidden layer in the encoder network was 128, the number of layers in the encoder network was two, the dropout rate for the encoder network was 0.1, the latent space had *K* = 10 dimensions, the latent distribution was standard normal, the gene likelihood was negative binomial (equivalently, a Poisson distribution with a gamma prior on the rate parameter as in Eq. (1)), and the dispersion pattern was constant per gene. For training, the batch size was 128, the learning rate was 5 × 10^−3^, early stopping was turned off, and the model was trained for 250 epochs on a NVIDIA 80 GB A100 GPU. We disabled early stopping to keep the training procedure consistent across LDVAE models and with Geneformer.

### Geneformer

To implement Geneformer [7], we used code from the its Hugging Face repository. This architecture incorporated the following: a BERT model type (bidirectional encoder representations from transformers), a maximum input size of 2048 tokens, six layers, four attention heads, an embedding dimension of 256, an intermediate size of 512 (twice the embedding size), and a rectified linear unit (ReLU) as the activation function. These models were trained on eight NVIDIA 32 GB V100 GPUs for 3 epochs with a linear learning scheduler, a learning rate of 0.001, and 10,000 warmup steps. Dropout rates were set uniformly at 0.02 for both attention and hidden layers. Lastly, the AdamW optimizer was used, alongside a layer norm epsilon of 1 × 10^−12^, an initializer range of 0.02, and a weight decay parameter of 0.001.

### Evaluation metrics

To assess the zero-shot performance of the LDVAE models, we evaluated their reconstruction accuracy for single-cell transcriptomic data. For each cell in an evaluation dataset, we computed the squared correlation *R*^2^ between the reconstructed and true gene expression values for the 1000 most highly variable genes (HVGs). Once again, let 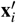 denote the reconstructed expression vector for the *i*-th cell. Within the LDVAE framework, this is modeled as 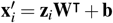 where, again, **z**_*i*_ is the latent representation for the *i*-th cell, **W** are the decoder weights, and **b** are the bias terms in the decoder. The Pearson correlation *r*_*i*_ between the true and reconstructed expression values for each cell is computed as the following

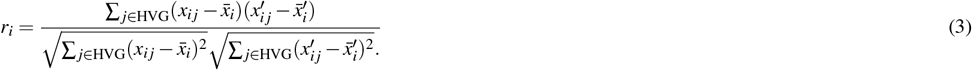

where 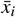 and 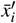 are the sample mean of the original and reconstructed gene expression for the *i*-th cell, respectively. For each cell, we calculate 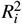 and summarize the performance across each dataset as the average reconstruction across all cells

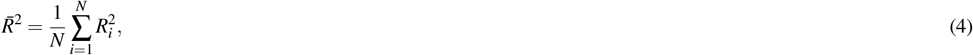

where *N* is the total number of cells in the evaluation dataset. This metric provides a single summary statistic for model performance.

To evaluate the zero-shot performance of the Geneformer models, we measured the Pearson correlation coefficient between the true and predicted rankings. Here, we follow Kedzierska et al. [31] and transformed the ordered outputs into rankings on the unit interval where the highest expressed genes were assigned a value of 1. The Geneformer software can output a sequence of up to 2,048 genes and, when input is passed in batches, the model outputs the sequence of the length equal to that of the longest input. Again following Kedzierska et al., in our evaluations, we limit the output sequence to the length of the input sequence.

To assess the robustness of our analyses for both LDVAE and Geneformer, we used paired t-tests on the *R*^2^ values of different models tested on the same sets of evaluation cells. The point of this exercise was to reflect the variability of predictions, essentially testing whether the *R*^2^ values for model A versus model B overlap on any evaluation. With the large number of cells per evaluation dataset, our analyses were highly powered, resulting in 224 of 225 model comparisons yielding significantly small standard errors. This is reflected in the *P*-values reported in **Table S1**. An example showing that model results were consistent across choices of random seeds can be seen, for example, in **Figure S1**.

### Assessing the similarity between models

For several follow-up analyses, we sought to estimate whether two different models provided similar representations of the same evaluation data. For example, in **Figure 3D**, we evaluated whether inclusion of malignant cells in training data changes model representations above and beyond what can be explained after training a model solely on healthy bone marrow data. To evaluate whether two models produce similar representations of an evaluation dataset, we used a linear regression-based framework. This approach provides a variable-wise estimate of whether the latent space of a “target” model can be explained by that of a “reference” model.

For a given pair of models, let 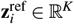 and 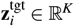 denote the reference and target, respectively, latent space representations of the *i*-th cell in an evaluation dataset, where *K* is the dimensionality of the latent space (assumed here to be identical for both models). For each *k*-th target latent variable, we fit a linear regression model using the reference latent variables as predictors

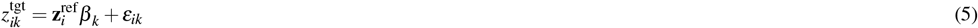

where *β*_*k*_ is a vector of regression coefficients and *ε*_*ik*_ is the residual error. Each regression model was trained on 70% of the evaluation dataset. To assess how well the *k*-th target latent variable is explained by the reference model, we computed the Pearson correlation *r*_*k*_ between the true 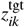 and the predicted values 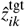 on the 30% held-out data. The *R*^2^ value for the *k*-th latent variable is then computed as the square of this correlation. Overall, this framework offers a detailed view on how well individual dimensions of the target latent space are able to be reconstructed from the reference model. Latent variables with low *R*^2^ scores indicate representations that are not well-explained by the reference model, highlighting potentially unique features in the training corpus used to generate the target representation (e.g., as described in **Figure 3I**). Our approach is related to existing methods such as canonical correlation analysis (CCA) and representational similarity analysis (RSA) [55]. However, whereas these methods provide a scalar summary of similarity across the entire latent space, our framework estimates similarity separately for each latent variable. This decomposition facilitates the identification of uniquely non-reconstructed dimensions in the latent space, offering a finer-grained analysis of model differences.

### Assessing how models generalize to unseen cell types

To better understand how the construction of a training corpus enables the ability of a model to generalize to unseen cell types, we trained the following variants of the Blood (Baseline) LDVAE approach. The details for these variations are given below.

- For the “No-Mono” model, we excluded the following cell types from the Blood (Baseline) training dataset (i.e., the healthy blood cells from scTab [25]) and used them for evaluation. These cells were annotated as “CD14-positive monocyte”; “classical monocyte”; “CD14-low, CD16-positive monocyte”; “CD14-positive, CD16-positive monocyte”; “monocyte”; “non-classical monocyte”; “CD14-positive, CD16-negative classical monocyte”; “macrophage”; and “intermediate monocyte”.
- For the “No-Bcell” model, we excluded the following cell types from the Blood (Baseline) training dataset (i.e., healthy blood cells from scTab [25]) and used them for evaluation. These cells were annotated as “naive B cell”; “B cell”; “memory B cell”; “transitional stage B cell”; “plasmablast”; “mature B cell”; “class-switched memory B cell”; “class switched memory B cell”; “plasma cell”; “immature B cell”; “pro-B cell”; “precursor B cell”; “IgA plasma cell”; and “IgG plasma cell”.

We computed reconstruction *R*^2^ for each cell as in Eqs. (3) and (4). To estimate reconstruction residuals, we subtracted the reconstructed cp10k expression values from the true cp10k expression values, averaged these differences across cells, and normalized by the mean cp10k expression of each gene.

### Assessing the impact of cancer training data

To better understand the impact of adding cancer cells into the training corpus, we trained (1) one baseline LDVAE with only healthy bone marrow data and (2) four LDVAEs trained with both healthy bone marrow data and the 3CA hematological subset described above but leaving out a different study. These studies included diffuse large B cell lymphoma cells from Steen et al. [44], multiple myeloma cells from Liu et al. [36], acute lymphoblastic leukemia cells from Caron et al. [37], and acute myeloid leukemia cells from van Galen et al. [50]. For each of these models, we computed reconstruction *R*^2^ values on the held-out cancer dataset. We additionally assessed the ability to reconstruct latent spaces of models trained on BoneMarrow+Cancer data from latent spaces of models trained only on BoneMarrow data as described above in Eq. (5).

### Assessing the impact of the transcription factor atlas on out-of-distribution performance

To extend our main set of analyses, we also evaluated each of our LDVAE models on two additional out-of-distribution datasets (see **Figure 3G**):

- **Heart**: For this evaluation, we trained a model where we held-out all cells with a tissue annotation labeled as “heart” in scTab [25]. To avoid contamination by hematopoietic cells, in the evaluation set we retained only the cell types with the following labels: “cardiac muscle cell”; “pericyte”; “capillary endothelial cell”; “fibroblast of cardiac tissue”; “endothelial cell of artery”; “smooth muscle cell”; “endothelial cell”; “vein endothelial cell”; “neuron”; “endothelial cell of lymphatic vessel”; “cardiac neuron”; and “mesothelial cell”. The final evaluation dataset consisted of 13,571 cells.
- **Kidney**: For this evaluation, we trained a model where we held-out all cells with a tissue annotation labeled as “kidney” in scTab [25]. To avoid contamination by hematopoietic cells, in the evaluation set we retained only the cell types with the following labels: “epithelial cell of proximal tubule”; “kidney loop of Henle thick ascending limb epithelial cell”; “endothelial cell”; “kidney collecting duct principal cell”; “kidney distal convoluted tubule epithelial cell”; “kidney interstitial fibroblast”; “kidney collecting duct intercalated cell”; “kidney connecting tubule epithelial cell”; “kidney loop of Henle thin descending limb epithelial cell”; “kidney loop of Henle thin ascending limb epithelial cell”; “renal interstitial pericyte”; and “vascular associated smooth muscle cell’. The final evaluation dataset consisted of 8,468 cells.

For our analysis of latent variables from the Blood+TFAtlas model, we estimated the ability to reconstruct latent representations of neurons in four “Blood+” models (i.e., Blood+BoneMarrow, Blood+BloodCancer, Blood+PerturbSeq, Blood+TFAtlas) from latent representations of neurons in the Blood (Baseline) model, as described above in Eq. (5). To estimate the correlation of model loadings with neuronal expression, we averaged cell-wise expression levels for each gene across all neurons in the Human Brain Cell Atlas evaluation dataset and correlated these expression means with factor-wise loadings (i.e., the columns of **W**) for each model.

## Data and code availability

All code is available under an open-source MIT license at https://github.com/microsoft/scFM-datamix. Specifically, we implemented LDVAE [18] which is implemented as part of `scvi-tools` [30] and downloaded from https://scvi-tools.org/ [54] (release 1.1.2). The version of Geneformer [7] we used corresponded to commit `b2bbd7ccc856e3b3c1a9199c7cca07d888d87663` (Jun 29, 2024) from the Hugging Face repository https://huggingface.co/ctheodoris/Geneformer/tree/main/geneformer. The Human Brain Cell Atlas (HBCA) [52] is publicly available online and can be accessed at github.com/linnarsson-lab/adult-human-brain. The acute myeloid leukemia (AML) dataset from van Galen et al. [50] and the squamous cell carcinoma (SCC) dataset from Ji et al. [51] are available in the Curated Cancer Cell Atlas (3CA) and can be found online at https://www.weizmann.ac.il/sites/3CA. Instructions for downloading the scTab corpus from the CELLxGENE census are provided at https://github.com/microsoft/scFM-datamix.

## Acknowledgments

This research was supported in part by a David & Lucile Packard Fellowship for Science and Engineering awarded to LC. SR acknowledges funding support from NCI K08 CA260442. Any opinions, findings, and conclusions or recommendations expressed in this material are those of the author(s) and do not necessarily reflect the views of any of the funders.

## Author contributions

AN, APA, and LC conceived the study. AN, AT, and MH developed the software and performed the analyses. AG and AWN conducted secondary analyses. NF, SR, PSW, APA, and LC provided resources. SR, PSW, APA, and LC supervised the project. AN, AT, APA, and LC wrote the initial draft. All authors interpreted the results and revised the manuscript.

## Supplementary Figures and Tables

**Figure S1.**
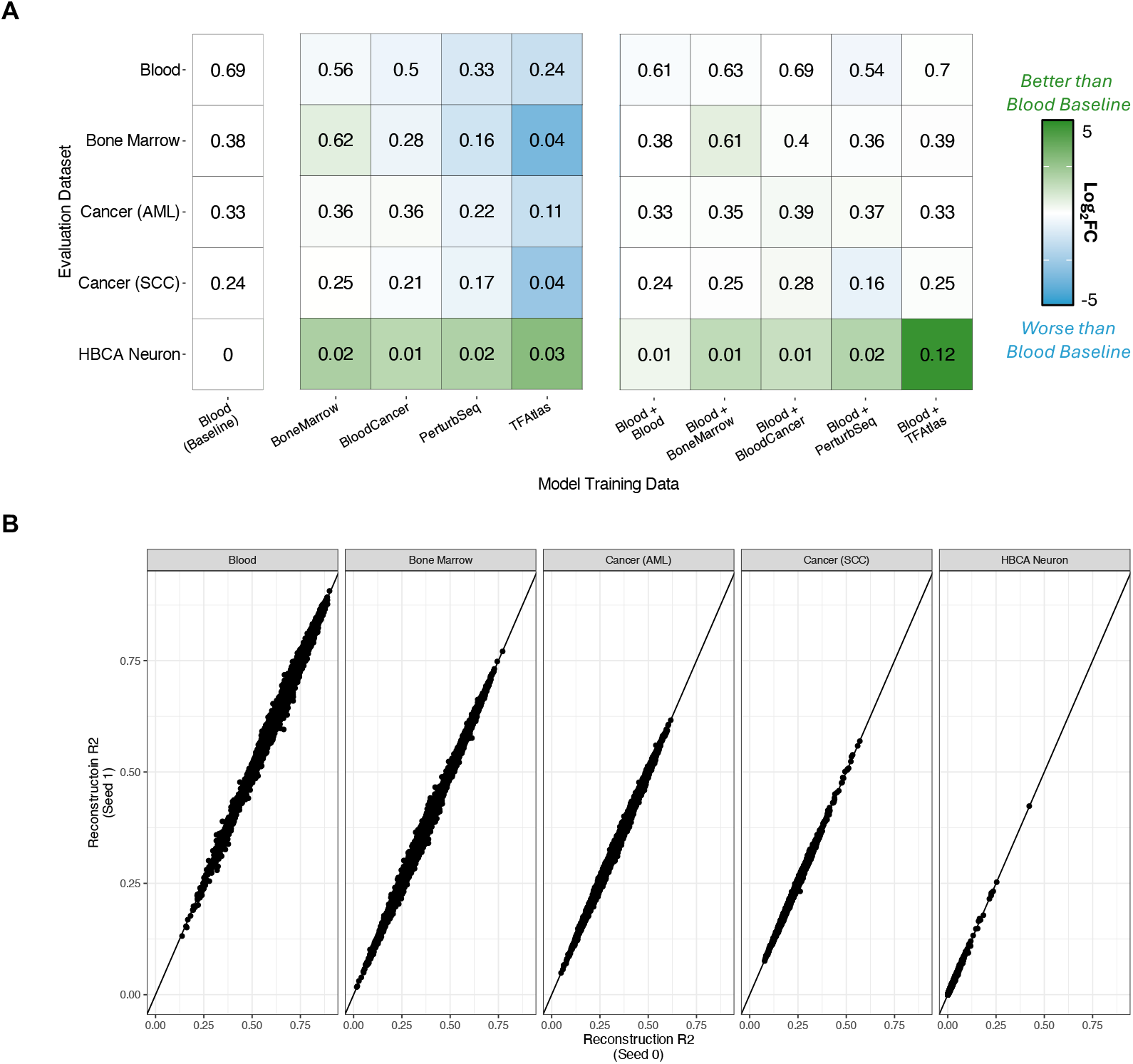
Performance of the linearly decoded variational autoencoder (LDVAE) models across different random seeds. **(A)** Analysis analogous to **Figure 1** but using a different random seed to create the training and evaluation datasets. The training data compositions are listed across each column. Here, the “Blood (Baseline)” model is trained exclusively on 100k healthy peripheral blood cells. The “BoneMarrow”, “BloodCancer”, “PerturbSeq”, and “TFAtlas” models are also trained on 100k cells coming exclusively from their respective studies. The “Blood+” combination models are all trained on 200k cells with 100k being healthy peripheral blood cells and the other 100k being from the other source. Numerical results in the heatmap are based on the average reconstruction accuracy (via *R*^2^) for highly-variable genes across each of the evaluation datasets shown across the rows. Within a given row, the color of each cell indicates the log_2_ fold change of the reconstruction *R*^2^ relative to the “Blood (Baseline)” model (with darker green representing greater relative performance for the other models). Note that all estimates are identical to the main analysis results up to two decimal points. **(B)** Plots comparing two different runs evaluating the cell-wise reconstruction *R*^2^ for the Blood (Baseline) model. Each point is a cell. See **Methods** for sample sizes.

**Figure S2.**
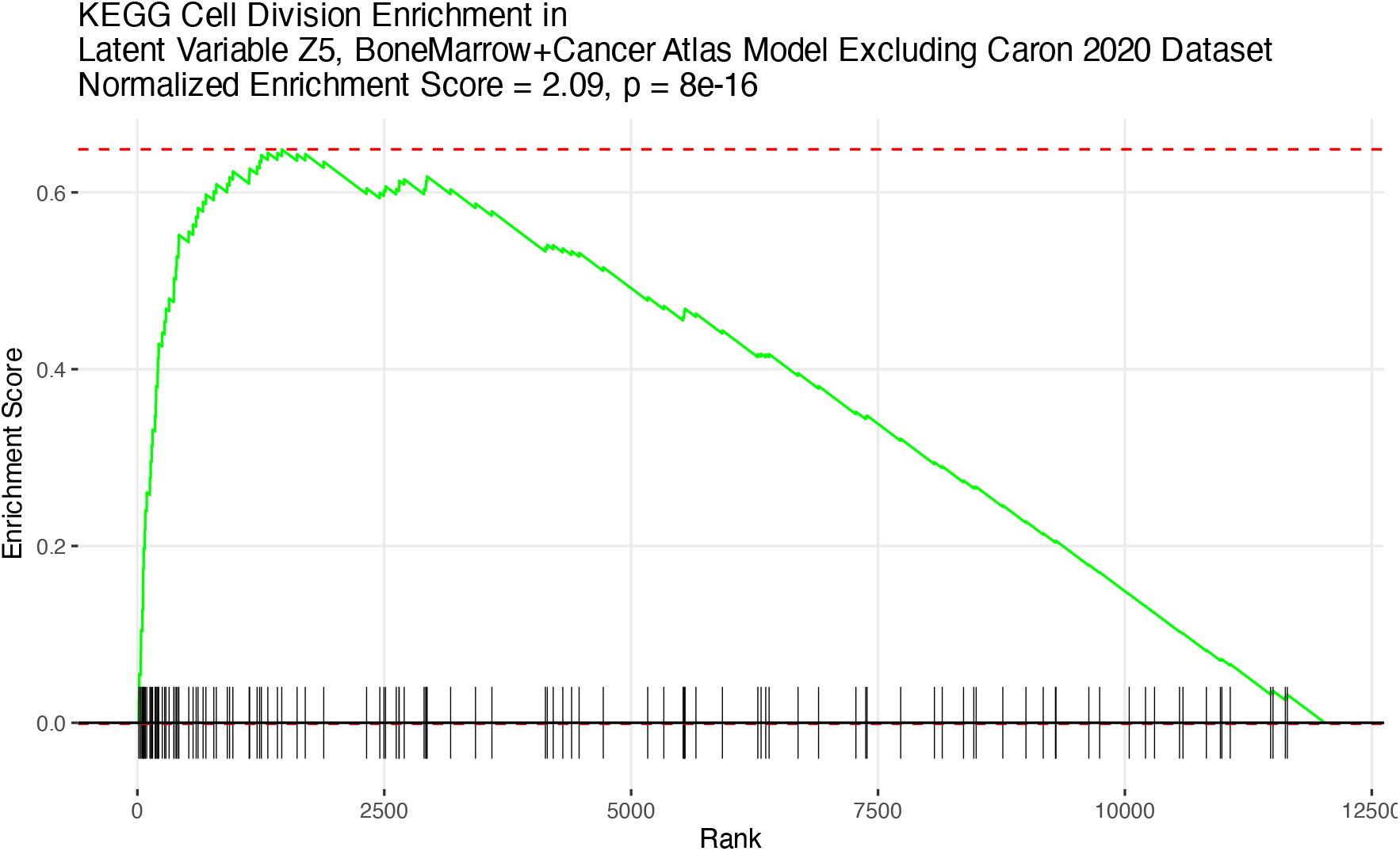
Gene set analysis shows an enrichment of mitosis in one of the latent variables learned by the BoneMarrow+Cancer model while holding-out the Caron et al. [37] acute lymphoblastic leukemia study. Depicted is a gene set enrichment analysis (GSEA) visualization for the KEGG (Kyoto Encyclopedia of Genes and Genomes) Cell Division gene set while analyzing the loadings on the latent variable *Z*_5_ from the BoneMarrow+Cancer model with the Caron et al. [37] acute lymphoblastic leukemia study held out. There are 124 genes in this gene set and 118 of them are included in our study, due to imperfect gene overlap between gene ontology and the presently analyzed datasets. GSEA yielded a normalized enrichment score of 2.09 and *P* = 8 × 10^−16^.

**Figure S3.**
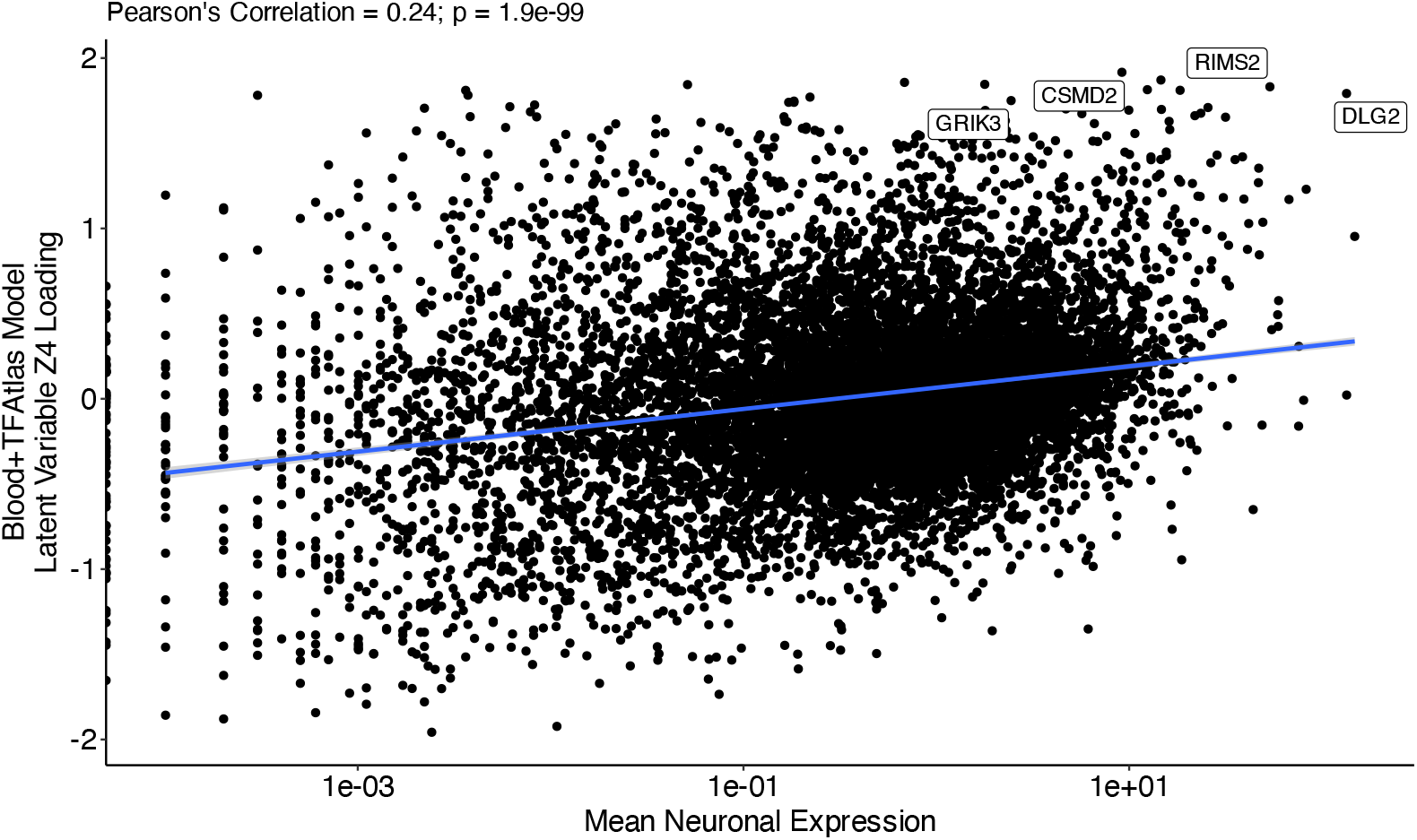
Correlation between the loadings on the latent variable *Z*_4_ learned by the Blood+TFAtlas model and neuronal gene expression. Here, the means of neuronal gene expression are derived using the Human Brain Cell Atlas (HBCA) evaluation dataset (**Methods**). This relationship has a Pearson correlation of 0.24 and *P* = 1 × 10^−99^, which was found by regressing the loadings from *Z*_4_ onto the log-mean-expression values with a linear model. Many of the top-loaded genes for this latent dimension are enriched for neuronal functions, especially synaptic function (e.g., *CSMD2, RIMS2*, and *DLG2*); one of the top-loaded genes is a subunit of the glutamatergic kaniate receptor, *GRIK3*.

**Figure S4.**
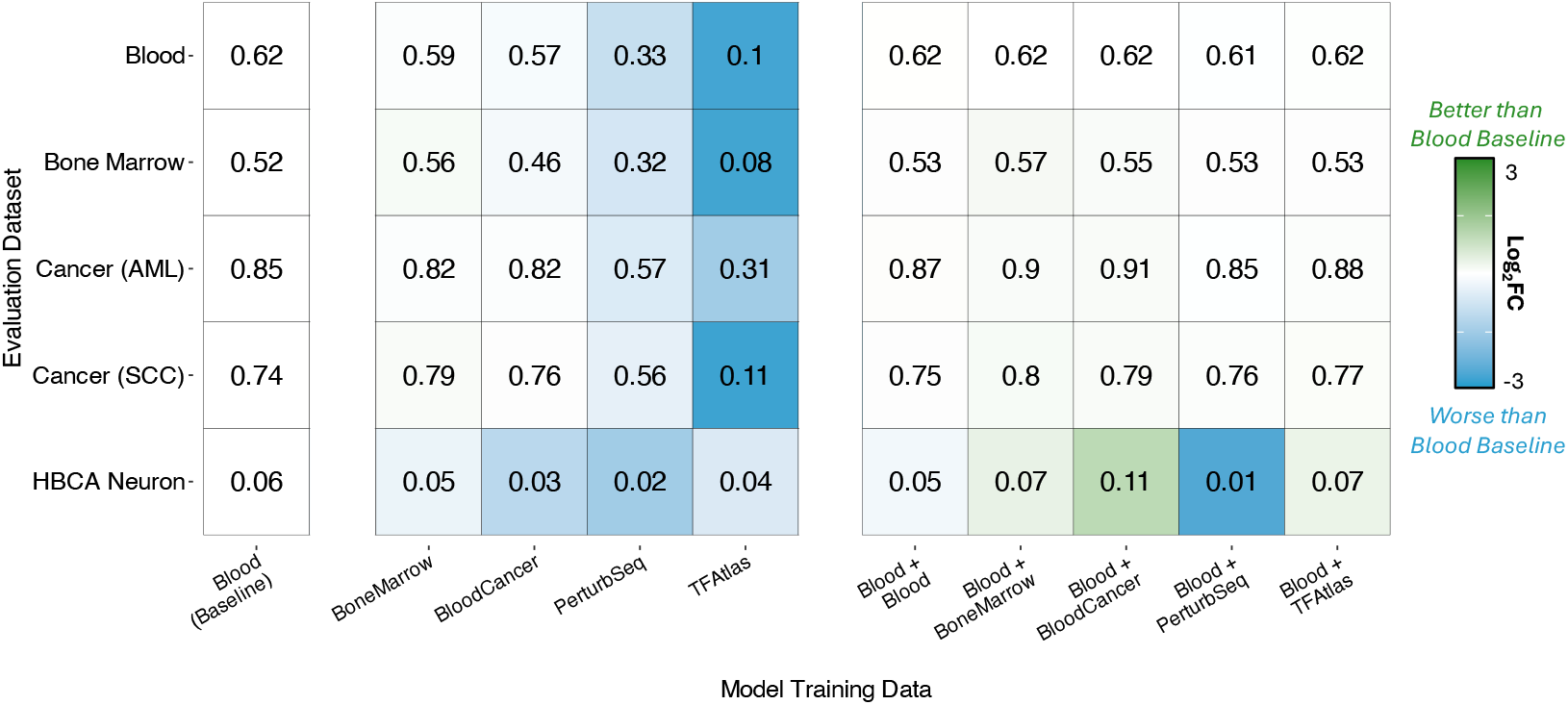
Estimates of reconstruction accuracy for various training and evaluation datasets using Geneformer. Different training data compositions are listed across each column. Here, the “Blood (Baseline)” model is trained exclusively on 100k healthy peripheral blood cells. The “BoneMarrow”, “BloodCancer”, “PerturbSeq”, and “TFAtlas” models are also trained on 100k cells coming exclusively from their respective studies. The “Blood+” combination models are all trained on 200k cells with 100k being healthy peripheral blood cells and the other 100k being from the other source. Since Geneformer produces ranked gene expression outputs rather than fully reconstructed expression estimates across all genes, numerical results in the heatmap are based on the Pearson correlation between the true and reconstructed gene expression ranks, averaged across cells and scaled to vary between 0 and 1 [31]. The evaluation datasets are shown across the rows. Within a given row, the color of each cell indicates the log_2_ fold change of correlation relative to the “Blood (Baseline)” Geneformer model (with darker green representing greater relative performance for the other models).

**Figure S5.**
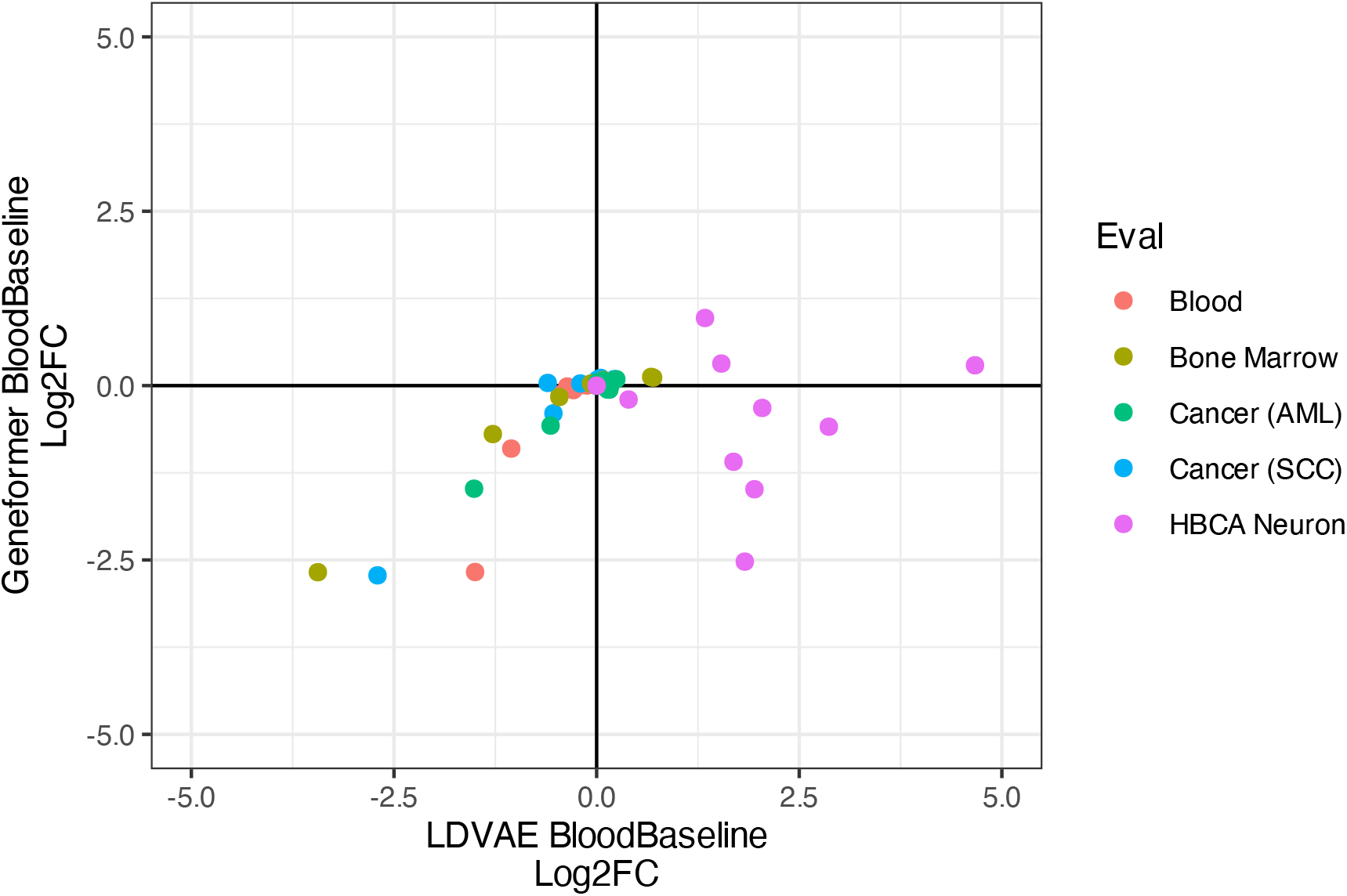
Comparing the performance of the LDVAE and Geneformer models using the Blood (Baseline). Each point shows the log_2_ fold change of the reconstruction *R*^2^ (LDVAE) and of the Pearson correlation of ranks (Geneformer) using the “Blood (Baseline)” model as a reference. Different points of the same color correspond to models with different training datasets.

**Figure S6.**
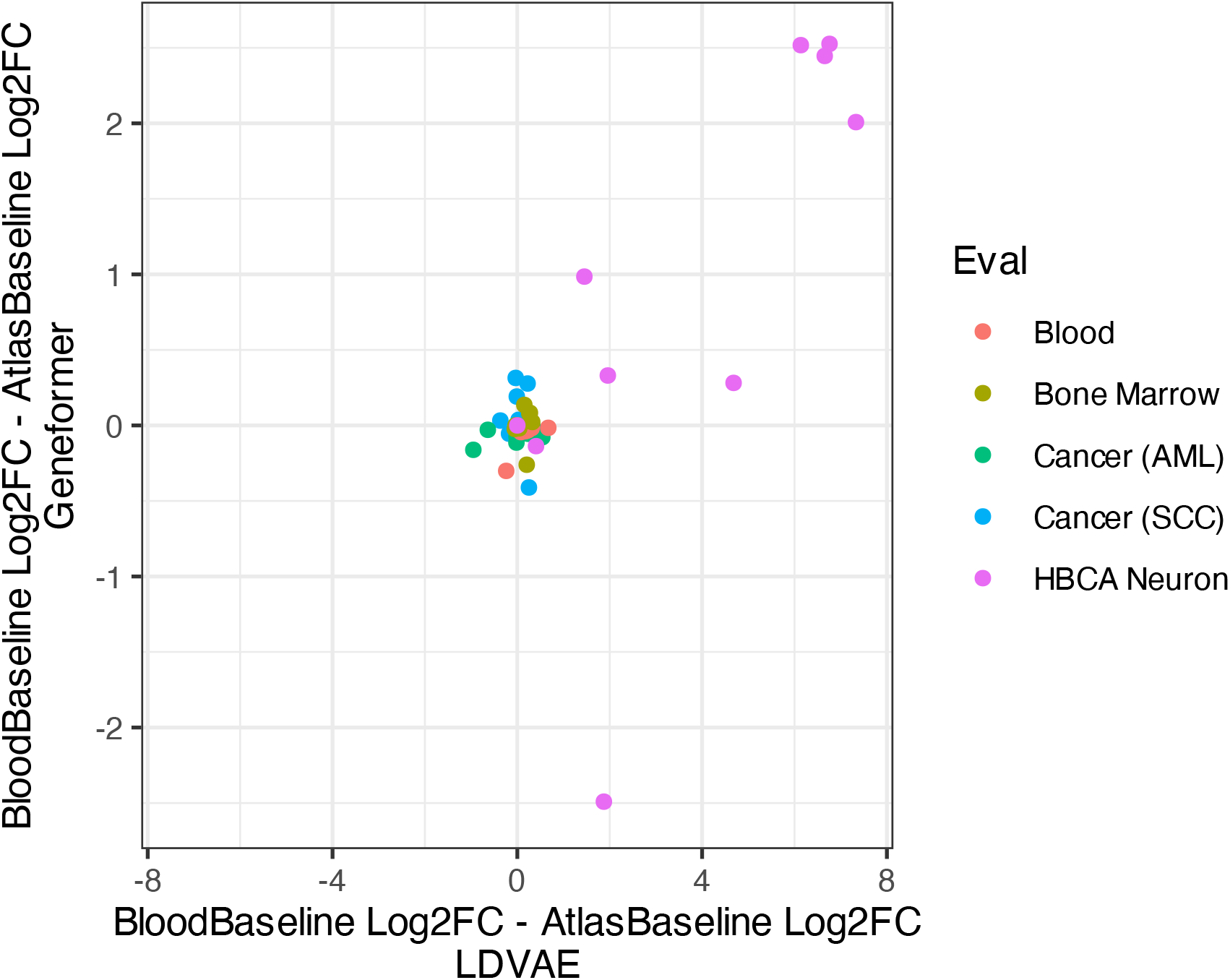
Performance of the LDVAE and Geneformer models when using the Blood (Baseline) versus the Atlas (Baseline). Each point shows the log_2_ fold change difference between the reconstruction *R*^2^ (LDVAE) and the Pearson correlation of ranks (Geneformer) using the “Blood (Baseline)” and “Atlas (Baseline)” models as references. Different points of the same color correspond to models with different training datasets.

**Figure S7.**
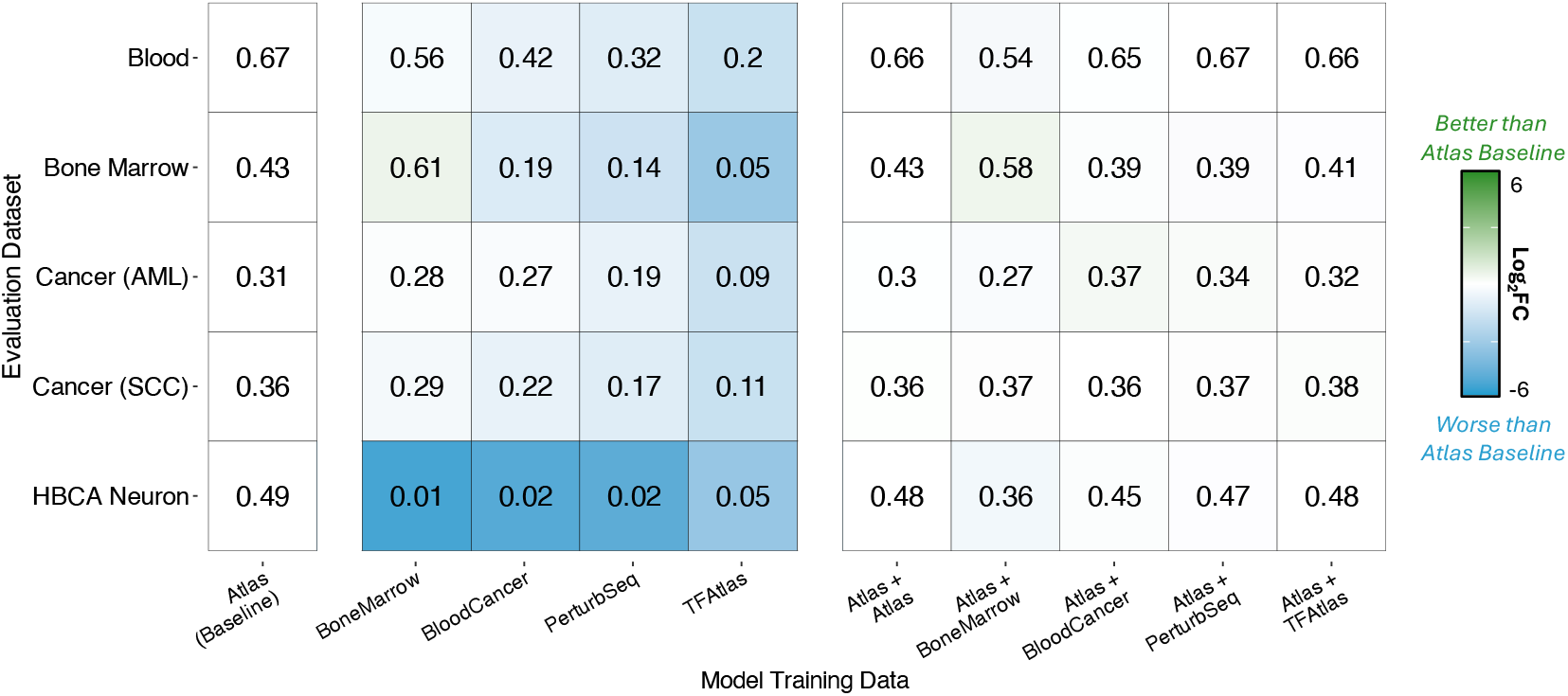
Estimates of reconstruction accuracy for various training and evaluation datasets using LDVAE and expanding the baseline training corpus to include a larger atlas. Different training data compositions are listed across each column. Here, the “Atlas (Baseline)” model is trained using a subsample of 100k cells taken from the entire scTab dataset. The “BoneMarrow”, “BloodCancer”, “PerturbSeq”, and “TFAtlas” models are trained on 100k cells coming exclusively from their respective studies. The “Atlas+” combination models are all trained on 200k cells with 100k being scTab and the other 100k being from the other source. Numerical results in the heatmap are based on the average reconstruction accuracy (via *R*^2^) for highly-variable genes across each of the evaluation datasets shown across the rows. Within a given row, the color of each cell indicates the log_2_ fold change of the reconstruction *R*^2^ relative to the “Atlas (Baseline)” model (with darker green representing greater relative performance for the other models).

**Figure S8.**
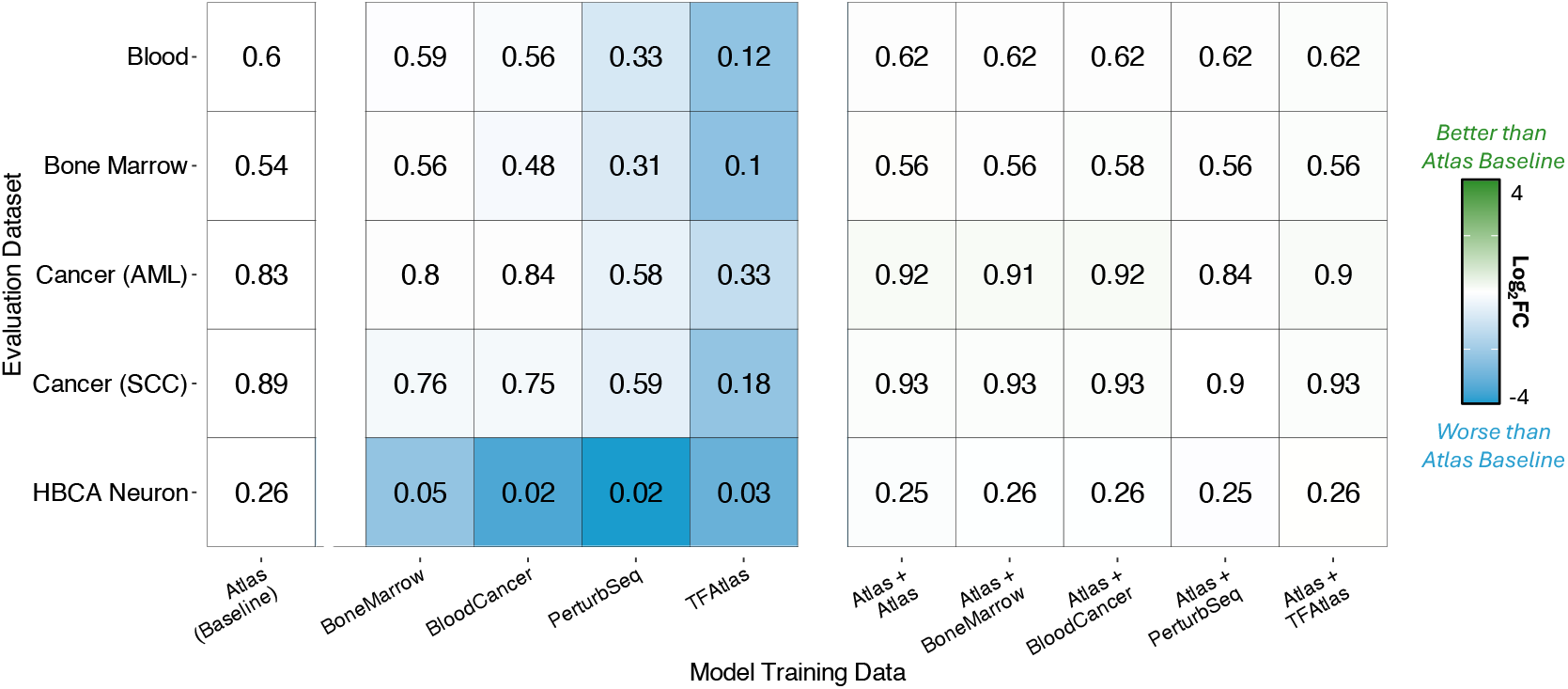
Estimates of reconstruction accuracy for various training and evaluation datasets using Geneformer and expanding the baseline training corpus to include a larger atlas. Different training data compositions are listed across each column. Here, the “Atlas (Baseline)” model is trained using a subsample of 100k cells taken from the entire scTab dataset. The “BoneMarrow”, “BloodCancer”, “PerturbSeq”, and “TFAtlas” models are trained on 100k cells coming exclusively from their respective studies. The “Atlas+” combination models are all trained on 200k cells with 100k being scTab and the other 100k being from the other source. Since Geneformer produces ranked gene expression outputs rather than fully reconstructed expression estimates across all genes, numerical results in the heatmap are based on the Pearson correlation between the true and reconstructed gene expression ranks, averaged across cells and scaled to vary between 0 and 1 [31]. The evaluation datasets are shown across the rows. Within a given row, the color of each cell indicates the log_2_ fold change of correlation relative to the “Atlas (Baseline)” Geneformer model (with darker green representing greater relative performance for the other models).

**Figure S9.**
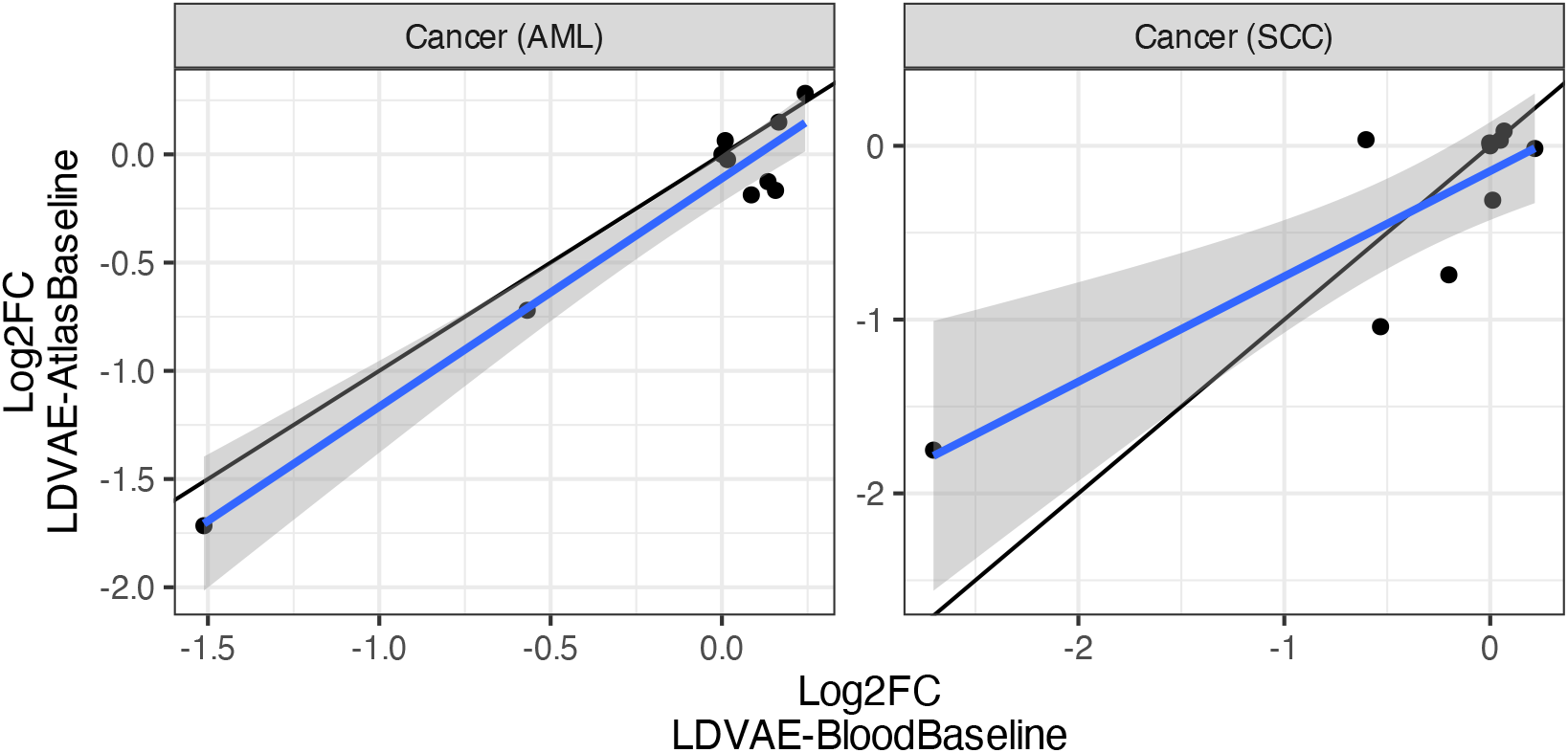
Performance of the LDVAE when using the Blood (Baseline) versus the Atlas (Baseline) on the cancer evaluation datasets, across all models visualized in Figure 2 and Figure S6. Each point shows the log_2_ fold change difference between the reconstruction *R*^2^ when using the “Blood (Baseline)” and “Atlas (Baseline)” models as references.

**Table S1. Assessing the robustness of predictive results across all experiments.** To assess the robustness of our analyses, we used paired t-tests on the *R*^2^ values of different models tested on the same sets of evaluation cells. The point of this exercise was to reflect the variability of predictions, essentially testing whether the *R*^2^ values for model A versus model B overlap on any evaluation. With the large number of cells per evaluation dataset, our analyses was highly powered, resulting in 224 of 225 model comparisons yielding significantly small standard errors and *P*-values (CSV).

